# Meiotic interactors of a mitotic gene *TAO3* revealed by functional analysis of its rare variant

**DOI:** 10.1101/033167

**Authors:** Saumya Gupta, Aparna Radhakrishnan, Rachana Nitin, Pandu Raharja-Liu, Gen Lin, Lars M. Steinmetz, Julien Gagneur, Himanshu Sinha

## Abstract

Studying the molecular consequences of rare genetic variants has the potential of identifying novel and hereto uncharacterized pathways causally contributing to phenotypic variation. Here we characterize the functional consequences of a rare coding variant of *TAO3*, previously reported to significantly contribute to sporulation efficiency variation in *Saccharomyces cerevisiae*. During mitosis *TAO3* interacts with *CBK1*, a conserved NDR kinase and a component of RAM network. The RAM network genes are involved in regulation cell separation and polarization. We demonstrate that the role of the rare allele *TAO3(4477C)* in meiosis is distinct from its role in mitosis by being independent of *ACE2*, which is a RAM network target gene. By quantitatively measuring cell morphological dynamics and conditionally expressing *TAO3(4477C)* allele during sporulation, we show that *TAO3* has an early role in meiosis. This early role of *TAO3* coincides with entry of cells into meiotic division. Time-resolved transcriptome analyses during early sporulation phase identified regulators of carbon and lipid metabolic pathways as candidate mediators. We experimentally show that during sporulation the *TAO3* allele genetically interacts with *ERT1* and *PIP2*, the regulators of tricarboxylic acid cycle and gluconeogenic enzymes, respectively. We thus uncover meiotic functions of *TAO3*, a mitotic gene and propose *ERT1* and *PIP2* as novel regulators of sporulation efficiency. Our results demonstrate that study of causal effects of genetic variation on the underlying molecular network has the potential to provide more extensive comprehension of the pathways driving a complex trait. This can help identify prospective personalized targets for intervention in complex diseases.

## INTRODUCTION

The ‘common disease, common variant’ rationale of genome-wide association studies is being challenged owing to limited fraction of disease heritability explained by the mapped common variants (Manolio *et al*. 2009; Zuk *et al*. 2014). One of the potential contributors to this ‘missing’ heritability has been suggested to be the potential effects of rare variants which are not considered (Saint Pierre and Génin 2014). This view has been substantiated by the identification of rare variants carrying a considerable risk for autism, schizophrenia and epilepsy (Stankiewicz and Lupski 2010). Thus characterizing the functional role of rare variants associated with complex diseases has the potential for revealing new biology and providing opportunities for treatment (Cirulli and Goldstein 2010; Zuk *et al*. 2014). Even though multiple variants for various diseases have been mapped, they have not been able to provide targets for treatment. This is because firstly, many variants have been mapped in regulatory or non-coding regions, therefore the affected gene is not known. Secondly, even when a variant is in a coding sequence but if the affected gene is not well characterized for the phenotype, the variant may be of little use. This makes characterization of either common or rare genetic variants laborious. Hence the causal path connecting a variant to the phenotype is usually unknown. Thus to understand the causal path of a variant, it is important to identify the mediating molecular pathways. Identifying these mediating pathways has the potential to greatly expand the set of possible targets for molecular intervention (Gagneur *et al*. 2013).

Yeast sporulation efficiency is a complex trait and many causative polymorphisms have been mapped in sporulation genes such as *IME1*, an initiator of meiosis (Gerke *et al*. 2009) and *RIM15*, a glucose-sensing regulator of meiosis (Lorenz and Cohen 2014). However a polymorphism each was identified in two genes with functional annotations described for mitotic growth only. These polymorphisms were in *MKT1*, a putative RNA-binding protein and *TAO3*, a putative scaffolding protein (Deutschbauer and Davis 2005). These novel non-synonymous polymorphisms, *MKT1(89G)* and *TAO3(4477C)* were identified in a high efficiency sporulating SK1 strain while the low efficiency S288c strain had *MKT1(89A)* and *TAO3(4477G)* (Deutschbauer and Davis 2005). In our previous work we determined that *MKT1(89G)* variant increased the sporulation efficiency by genetically interacting with regulators of mitochondrial retrograde signaling and nitrogen starvation during sporulation (Gupta *et al*. 2015). Tao3 encodes a highly conserved scaffolding protein that is a component of the RAM (Regulation of Ace2p activity and cellular Morphogenesis) signaling network. Tao3 is required for activation and localization of an NDR protein kinase Cbk1, another essential component of RAM network (Du and Novick 2002; Hergovich *et al*. 2006). This RAM signaling network including Cbk1, Hym1, Kic1, Mob2 and Tao3 contributes to various important processes of mitotic cellular growth. Regulation of transcription factor Ace2 by the RAM network is critical for cell separation and polarized growth (Nelson *et al*. 2003). Ace2 peaks early in mitosis and is involved in G_1_/S transition (Spellman *et al*. 1998). This RAM network also known to regulate cellular progression through a Ace2-independent pathway (Bogomolnaya *et al*. 2006). However none of these *TAO3* mitotic interactions provide clues for its role in the developmental process of meiosis and sporulation.

Here we characterized the functional role of *TAO3(4477C)* in sporulation efficiency variation by elucidating the molecular pathways linking this mitotic gene in meiosis. We compared phenotypes of a pair of S288c-background strains differing only for the casual *TAO3* polymorphism. Genome-wide transcriptional dynamics during sporulation of these strains identified candidate mediator genes. Allele-specific genetic interaction assay between these candidate genes and casual *TAO3* allele identified the regulators of tricarboxylic acid cycle and gluconeogenic enzymes as casual and novel regulators of sporulation efficiency.

## RESULTS

### Role of causative allele of *TAO3* in sporulation efficiency variation

Analysis of *TAO3* sequence of 38 *S. cerevisiae* strains in the Saccharomyces Genome Resequencing Project (SGRP) database (Liti *et al*. 2009) and 24 strains in Saccharomyces Genome Database (SGD), showed that *TAO3(4477C)* allele of SK1 strain was a rare variant (minor allele frequency = 1.6%, Figure 1A). Deutschbauer and Davis (2005) mapped this rare variant as a causal allele for increased sporulation efficiency in a cross between S288c (low sporulating) and SK1 (high sporulating) strains and introduced it in the S288c background to construct allele replacement strain YAD331. This allele replacement strain differed from the parental S288c strain (*TAO3(4477G)*) only for this variant. A clean allele replacement strain, termed as “T strain” was constructed from YAD331 (see Methods) and its sporulation efficiency at 48h was reconfirmed to be three-fold higher than the S288c strain (termed as “S strain”, *P* = 1.8 × 10^−10^, pair test in Methods, Figure 1B). This fold-difference remained constant even after a week of incubation (Figure 1C). Studying the progression of meiotic phases showed that the T strain initiated meiosis within 12h (Figure 1D–E). Quantitative comparison of the time to initiate meiosis and the rate of transition from G_1_/G_0_ into Meiosis I stage showed significant difference between the T and S strains (Figure 1D–E, Figure S1). This suggested that *TAO3(4477C)* affected entry of the T strain cells initiating meiosis within 12h in sporulation. To resolve when during this 12h time phase *TAO3(4477C)* affects the phenotype, this endogenous allele in the T strain was placed under a tetracycline-responsive promoter (P_Tet_-*TAO3(4477C)* strain, see Methods). In the absence of tetracycline analogue, doxycycline, P_Tet_-*TAO3(4477C)* strain showed higher expression of *TAO3(4477C)* relative to its expression in the S strain (Methods, Figure S2). Addition of 2μg/ml doxycycline significantly reduced *TAO3* expression level, making it equivalent to the S strain (Methods, Figure S2). Concomitantly in the absence of doxycycline, the P_Tet_-*TAO3(4477C)* strain showed high sporulation efficiency (Figure 1F). In presence of doxycycline for the entire 48h in sporulation medium, the sporulation efficiency of the P_Tet_-*TAO3(4477C)* strain was equivalent to the S strain (Figure 1F). This suggested that high *TAO3(4477C)* expression was required for the high sporulation efficiency phenotype. We next reduced *TAO3(4477C)* expression for specific shorter time-periods in the sporulation medium. Sporulation efficiency of the P_Tet_-*TAO3(4477C)* strain was equivalent whether doxycycline was present only for the first 6h or for 48h and this efficiency was equivalent to the S strain (Figure 1F). However when doxycycline was present only for the first 3h, the P_Tet_-*TAO3(4477C)* strain showed a slight but significant difference (*P* = 0.02) in sporulation efficiency compared to S strain (Figure 1F). This showed that *TAO3(4477C)* allele affected sporulation efficiency within the first 6h in sporulation.

**Figure 1.**
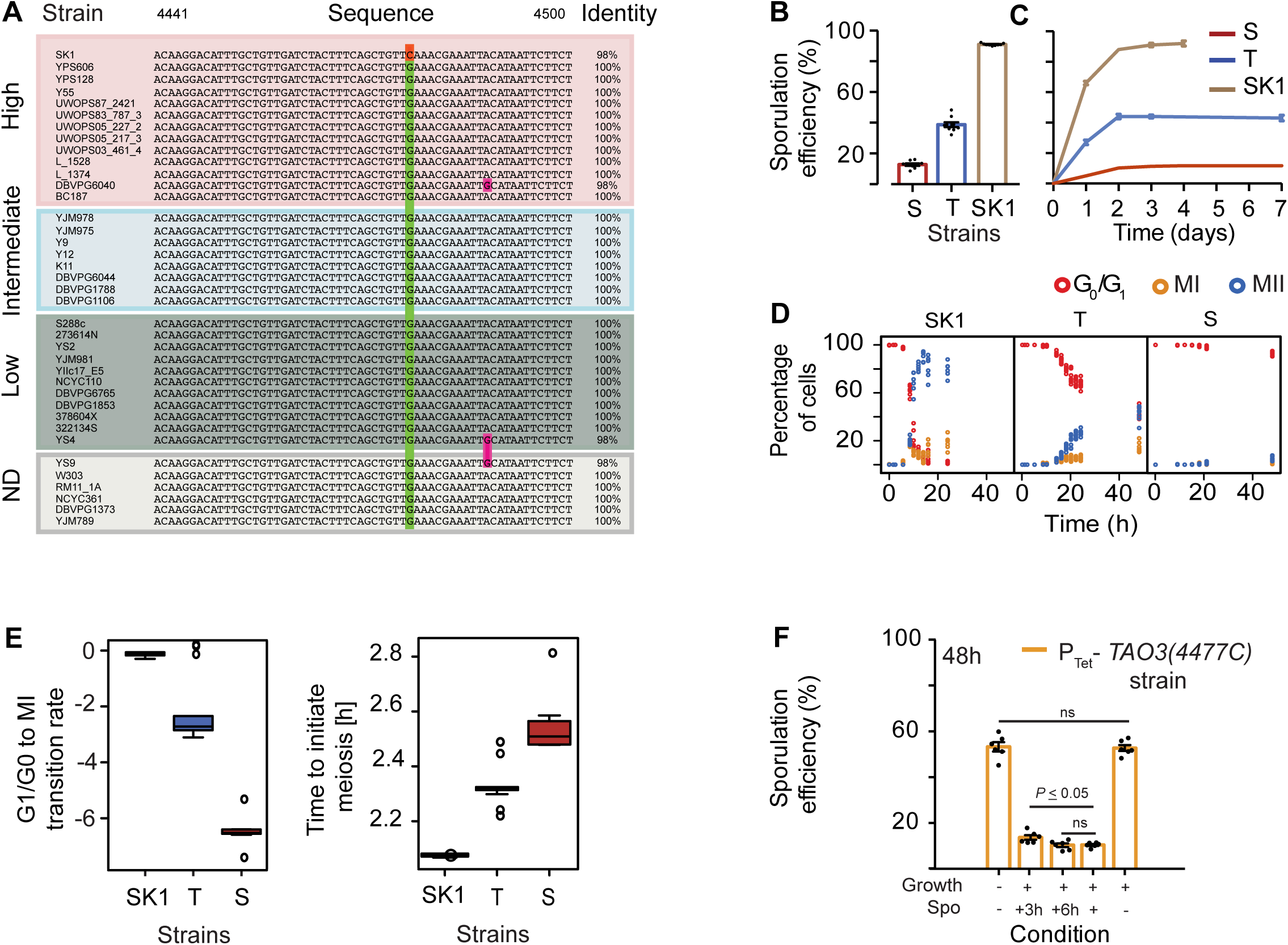
Role of *TAO3* in sporulation efficiency. (A) Comparison of genomic sequence of *TAO3* (4,441-4,500) across the SGRP collection (Liti *et al*. 2009). The 4,477th position of *TAO3* consists of the sporulation causative variant where identical nucleotides are indicated by the same color. Identity indicates the percentage match between the nucleotides in the shown region of the gene. The strains are ordered according to their mean sporulation efficiency (Tomar *et al*. 2013): high (60-100%), intermediate (10-60%), low (0-10%) and ND (not determined). (B) Bar plots represents the mean sporulation efficiency after 48h of the SK1, T and S strains. The sporulation efficiency data is indicated as circles. (C) Line graphs represent the mean sporulation efficiency of the S, T and SK1 strains measured till saturation, i.e. till sporulation efficiency did not vary for 3 consecutive days. (D) Percentage of 1-, 2- and 4-nuclei states of the T strain (y-axis) versus time in sporulation medium (x-axis). 1-nucleus stage is indicated as red circles (G_0_/G_1_ phase), 2-nuclei state as yellow circles (completion of Meiosis I, MI phase) and blue circles is 4-nuclei stage (completion of Meiosis II, MII phase). (E) Bootstrap distribution of the time to initiate meiosis and the rate of transition from G_1_/G_0_ into MI, estimated from time courses in (D). See Methods for details. (F) Conditional expression of *TAO3(4477C)* during sporulation in P_Tet_-*TAO3(4477C)* strain (indicated as P_Te_t). Y-axis is the mean sporulation efficiency in 48h. No doxycycline in growth (YPD) or spo (YPA + sporulation) medium is depicted as “-” condition on x-axis and addition of doxycycline is depicted as “+” in that condition. “+3h” condition in Spo implies doxycycline was throughout in the growth medium and in the sporulation medium till 3h after which cells were sporulated in the absence of doxycycline. “+6h” condition implies doxycycline was throughout in the growth medium and in the sporulation medium till 6h after which cells were sporulated in the absence of doxycycline. *P* value was calculated by an unpaired t-test. Error bars are standard error of mean.

### Role of *TAO3* in meiosis is distinct from its role during mitosis

Varying the gene expression of *TAO3(4477C)* affected the sporulation efficiency phenotype. Hence to identify the molecular pathways affected by this causative allele, we studied the global gene expression dynamics during sporulation in the allele replacement strains. Time-resolved transcriptomics of the T and S strains were compared from 0h to 8h30m in sporulation medium (see Methods). At the initial time point (t = 0h) only 190 out of 6,960 transcripts (~3%) showed differential expression, with an enrichment for a single gene ontology term iron ion homeostasis (*P* = 0.04, post Holm-Bonferroni corrected, Figure S3). In contrast 1,122 transcripts (including non-coding SUTs, Table S1) showed statistically significant differences in gene expression dynamics as a function of time between the two strains (FDR cut-off 10%, when controlling for expression at t = 0h). While *TAO3* was amongst the transcripts showing differential expression dynamics during sporulation (*P* = 0.004), none of its mitotic interactors showed differential expression (Figure 2B). However a few *ACE2*-regulated genes did show differential expression (11 genes labeled green in Figure 2B), so we studied the effect of *ace2*Δ in the T strain and high sporulating SK1 strain. *ACE2* is known to regulate the budding phenotype (Voth *et al*. 2005) thus both the T and SK1 strains with *ace2t*Δ showed clumping. However *ace2*Δ did not affect sporulation efficiency of either the T or SK1 strain (Figure 2C). Ace2-independent effect of RAM network on cellular polarization have been observed previously (Nelson *et al*. 2003) therefore it is possible that this network could still be involved in meiosis.

To determine whether the mitotic interactors of *TAO3* were distinct from its meiotic interactors, we again used P_Tet_-*TAO3(4477C)* strain and reduced *TAO3* expression only during the mitotic growth phase, i.e. in glucose rich (YPD) medium. We observed no growth difference between P_Tet_-*TAO3(4477C)* strain with or without doxycycline and the T strain (Figure S2). In addition there was no effect on sporulation efficiency among the strains (Figure 1F). These results implied that probably *TAO3(4477C)* allele had a distinct meiotic role from its established function in mitosis-related processes.

**Figure 2.**
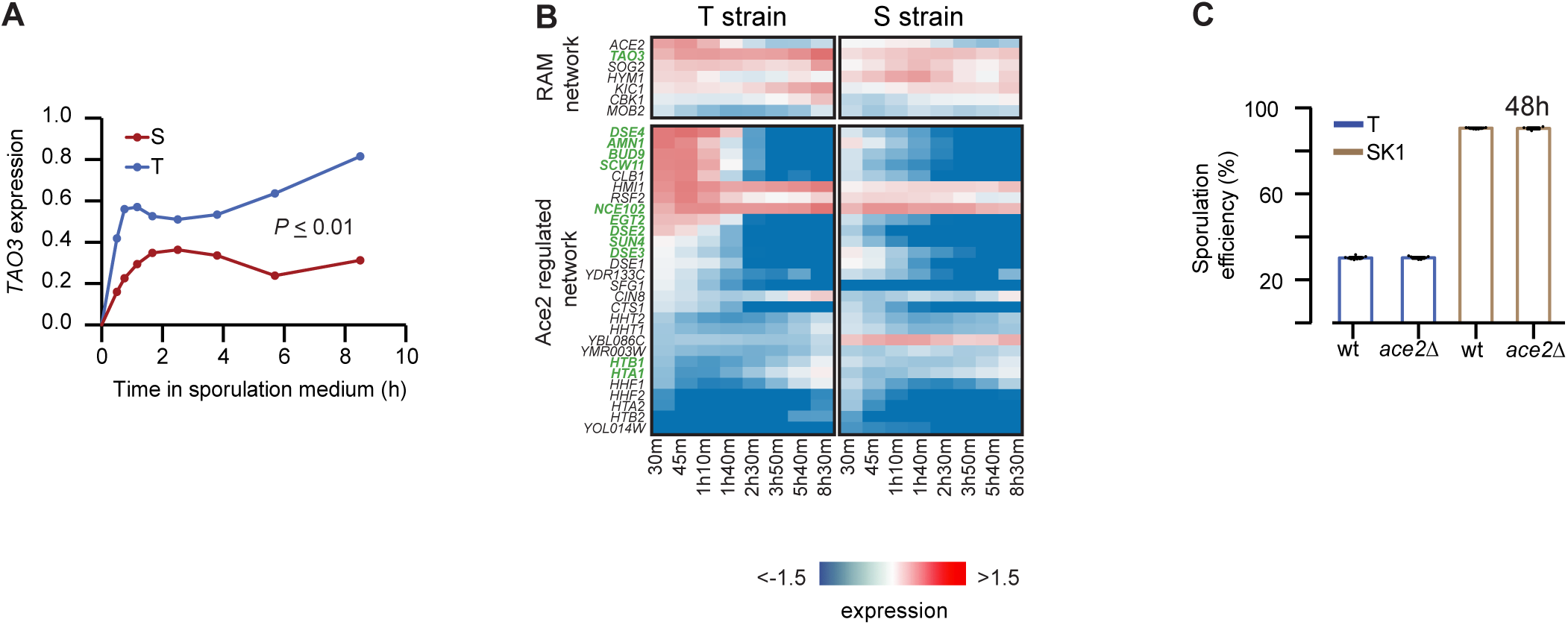
Role of *TAO3* in meiosis is distinct from its role during mitosis. (A) Heatmap showing gene expression of RAM network genes and Ace2-regulated genes in the T and S strains. Gene names in green show differential expression (data in Tables S1 and S11). (B) Bar plots represent the mean sporulation efficiency after 48h of the SK1 and T wild type (wt) and *ace2*Δ deletion strains. Pair and interaction tests (described in Methods) were performed to test significance. (C) Expression profile (log_2_ fold change t_0_) of *TAO3* is given in the y-axis for the T (purple) and S strains (red) and the x-axis denotes the time in sporulation medium (data in Tables S1 and S11).

### Temporal gene expression profiling predicts *TAO3(4477C)*-specific interactors during sporulation

We showed that *TAO3(4477C)* had a distinct role in the sporulation processes within first 6h of sporulation (Figure 1F). Hence we identified by clustering (see Methods), sets of differentially expressed genes showing early and increasing trend in their expression profiles in the T strain only. Various sporulation genes including crucial regulators of meiosis, namely *IME1, IME2, DMC1* and *NDT80* were enriched (*P* = 5.5 × 10^−12^) in a cluster showing increasing expression (Cluster II) during sporulation in the T strain (Figure 3B, see Methods). Approximately 50% of Cluster II genes of the T strain showed a similar increasing trend in the S strain, including *IME1, IME2, DMC1, ECM11* and *NDT80* (Figure S4, Table S2). Interestingly very few early expressing genes (Cluster I) of the T strain overlapped with the S (7%, Figure S4). These genes belonged to biological processes that regulated entry into sporulation, such as carbohydrate metabolic process, ion transport, mitochondrial organization and cellular respiration (Table 1). Furthermore genes involved in biological processes like carbohydrate metabolic process and mitochondrial organization showed repression in the S strain (Table 1, Figure S5). Therefore to study the early effects of the causal *TAO3* allele, we identified regulators of only those differentially expressed genes that showed early and increasing expression uniquely in the T strain (Tables S3 and S4).

**Figure 3.**
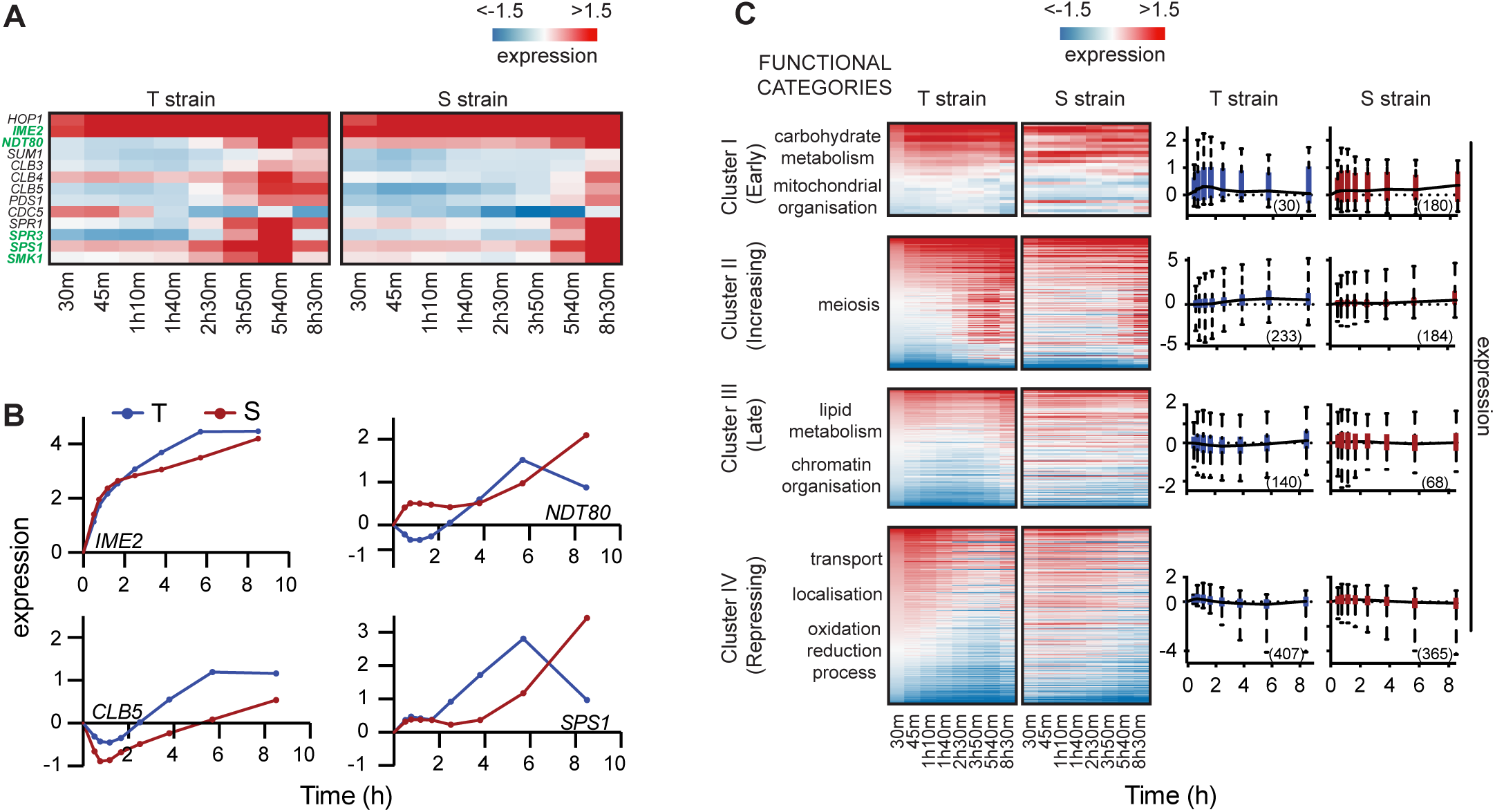
Global gene expression variation in presence of causative *TAO3* allele. (A) Temporal heat map of meiotic genes in the T and S strains. The gene names shown in green are differentially expressed in the presence of *TAO3(4477C)*. (B) The expression profile (log_2_ fold change t_0_) for the meiotic landmark genes is given in the y-axis and the x-axis denotes the time in sporulation medium. Red line represents the expression profile of the respective gene in the S strain and blue line is the same in the T strain. (C) Heat map of the T and S strains showing differentially expressed gene across time within each cluster. Each row represents a single gene and columns are time points of each strain (for gene list in each cluster see Table S2). The order of genes in the two strains is based on the clustering of the T strain. Functional GO categories of genes in each cluster are shown on left. The boxplots shown on right represent the average expression profile of each cluster in the T and S strains. The number of genes in each cluster in a strain is indicated in brackets.

**Table 1.**
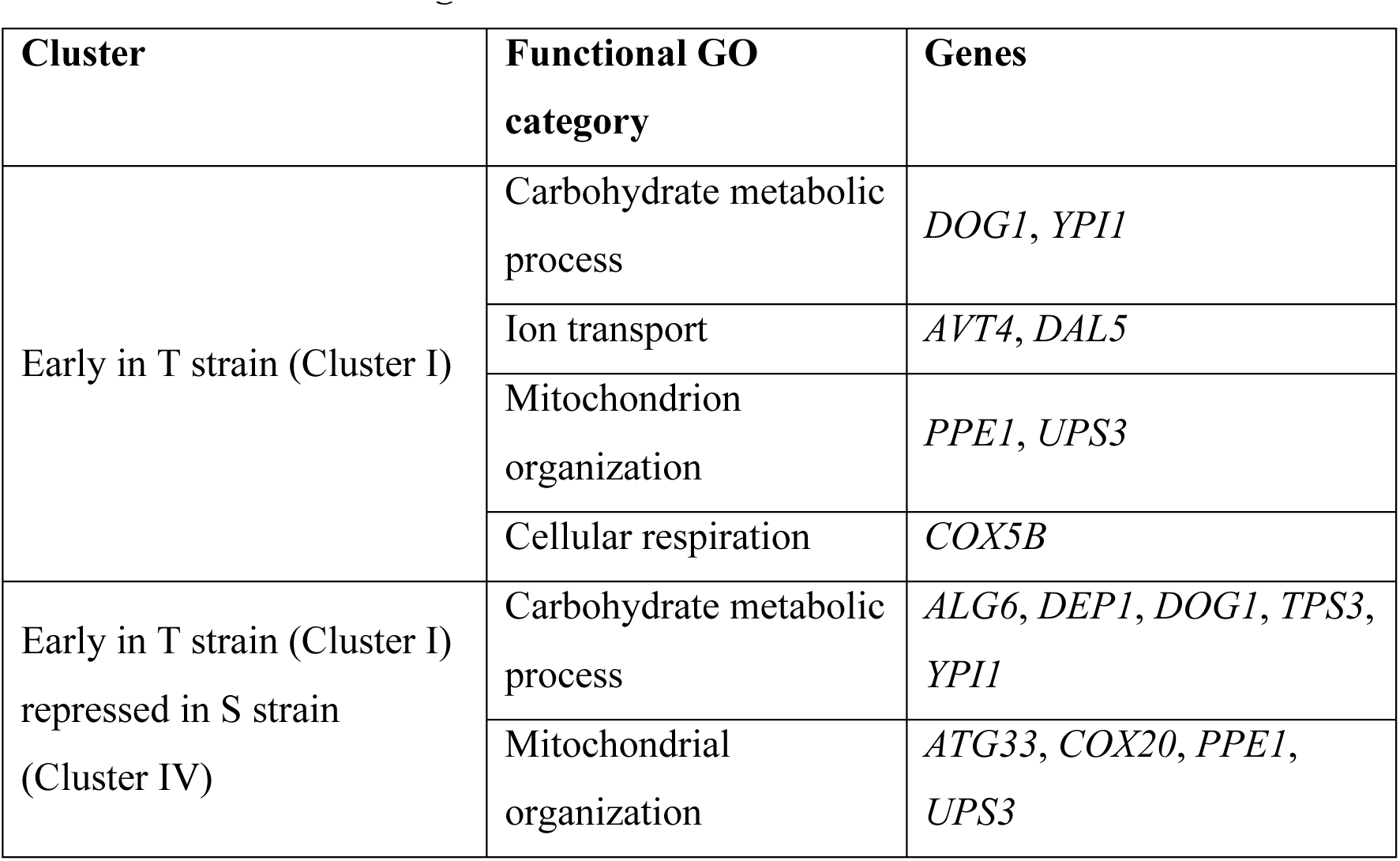
Functional GO categories of clusters in the T and S strains.

These regulators were enriched in nutrient metabolism and chromatin modification - biological processes important for initiation of meiosis (Neiman 2011). A core sporulation gene *UME6*, which together with *IME1* induces expression of early meiotic genes (Kassir *et al*. 2003) and is known to regulate other important processes for initiating meiosis (Table 2, Lardenois *et al*. 2015) was also identified. Interestingly among the regulators of the genes showing early and increasing expression uniquely in the T strain, we identified 25 upstream regulators of *UME6* (Figure 4A, Tables S3-5). Among these regulators were *ERT1, OAF1-PIP2* and *DAL81. ERT1*, a regulator of carbon source utilization (Turcotte *et al*. 2010) is involved in the switch from fermentation to respiration in glucose-limiting conditions (Gasmi *et al*. 2014). *OAF1-PIP2* is a protein complex regulating lipid metabolism (Karpichev and Small 1998). *DAL81* is a regulator of nitrogen degradation pathway (Marzluf 1997). Interestingly like *UME6*, *OAF1* target genes were repressed in the S strain (Cluster IV, Table S6). Earlier work in S288c and SK1 strains has shown upregulation of *ERT1, PIP2* and *DAL81* in SK1 strain during sporulation (Primig *et al*. 2000). However their deletion in S288c strain had no effect on its sporulation efficiency (Deutschbauer *et al*. 2002). A few other interesting candidate genes that were not upstream *UME6* were also identified (Tables S3 and S4). These included *GAT1*, a regulator of nitrogen metabolism (Ljungdahl and Daignan-Fornier 2012) and *GAT3*, a regulator of spore wall assembly (Lin *et al*. 2013). We next tested if the metabolic regulators identified through this analysis were *TAO3(4477C)-* specific mediating genes during sporulation.

**Table 2.**
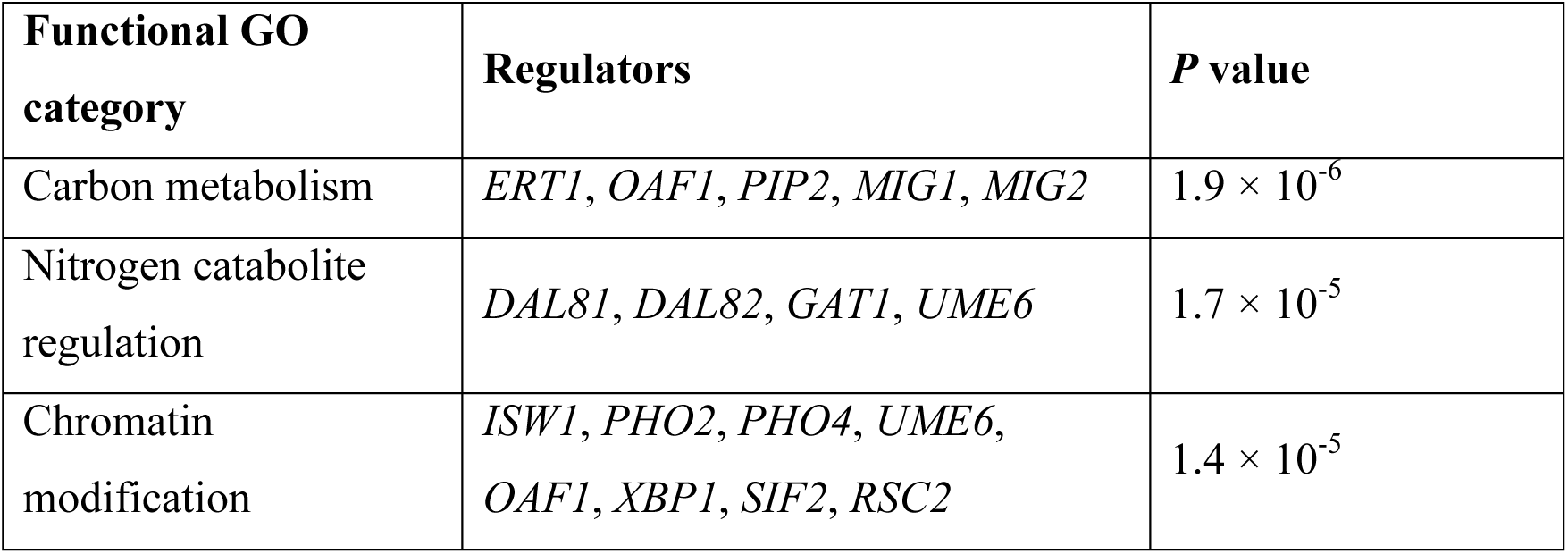
Functional GO categories of regulators of clusters in the T and S strains.

**Figure 4.**
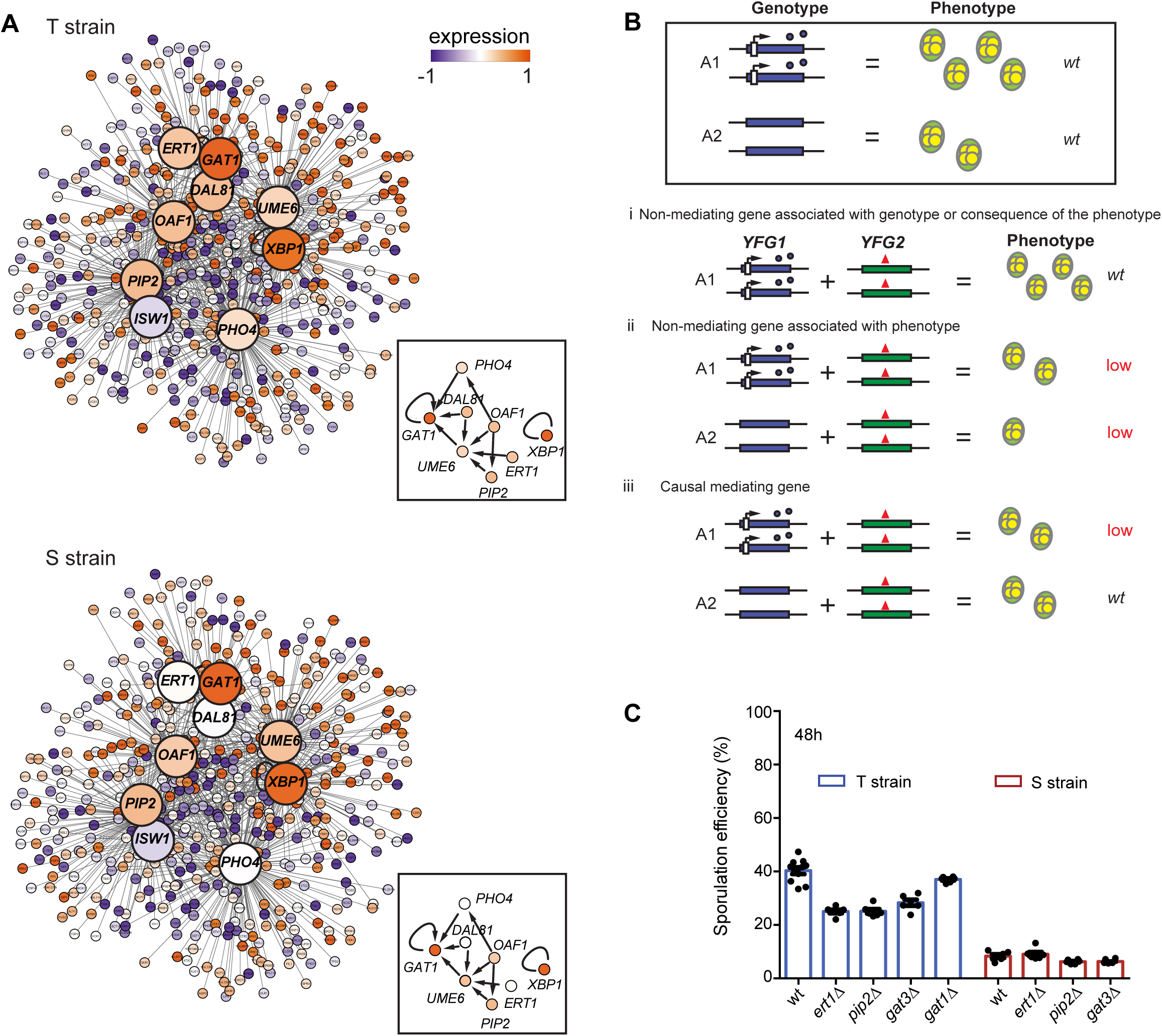
Identifying candidate genes mediating the allele specific effects of *TAO3* during sporulation using the temporal gene expression data. (A) Regulatory network of candidate genes predicted to mediate the effects of *TAO3(4477C)* in sporulation. The candidate mediating genes are shown as bigger nodes (large circles) with their target genes (small circles) connected to them as straight lines. The box contains the protein network interactions of the candidate genes with the core sporulation gene *UME6*, obtained from YEASTRACT (see Methods). Colors inside the nodes were calculated as an average of the first six time points in sporulation (early phase). For complete list of interacting genes and their expression values, see Tables S5 and S11, respectively. (B) Genetic model for functional validation of allele-specific interactors mediating sporulation efficiency variation. Wild type effect comparison of the two alleles A1 and A2 of *YFG1* gene is shown inside the box. A1 is associated with high sporulation efficiency (wild type genotype and phenotype shown) and A2 is associated with low sporulation efficiency (wild type genotype and phenotype shown). Genetic interaction of these *YFG1* alleles with candidate mediating genes (*YFG2*) is represented - (i) Representation of non-mediating gene associated with genotype only or is a consequence of the phenotype since *yfg2*Δ in the presence of A1 does not affect the wild type phenotype of A1; (ii) Representation of nonmediating gene associated with the phenotype independent of the allele since *yfg2*Δ in the presence of both A1 and A2 lowers (low) the phenotype; (iii) Representation of causal mediating gene since *yfg2*Δ only in presence of allele A1 lowers the phenotype and in the presence of allele A2 does not change the wild type phenotype of A2. (C) Bar plots represent the mean sporulation efficiency after 48h of the T and S wild type (wt) and *ertlΔ, pip2*Δ and *gat3*Δ strains. Pair and interaction tests (see Methods) were performed to test significance.

### Allele-specific functional validation identifies *TAO3(4477C)*-specific genetic interactors during sporulation

The candidate genes predicted in the above analysis could be either causal mediating genes interacting with *TAO3(4477C)* during sporulation or non-mediating consequential genes associated with only the genotype or the phenotype. To identify only the causal mediating genes, we used a genetic model described previously (Figure 4B, Gupta *et al*. 2015). According to this model if a gene is associated only with the genotype and not with the phenotype or is expressed as a consequence of the phenotype, its deletion would not affect the T strain phenotype. If a gene had an independent role in sporulation phenotype, its deletion will result in both a reduction in phenotype and an additive effect, irrespective of the genetic background. Any significant deviation from this expectation would imply dependence on the genotype with epistasis being an extreme case. In this scenario deleting the gene in T strain would affect the phenotype while deleting the same in the S would not have an effect on the phenotype, making it a causal mediating gene. While *gat1*Δ had no effect on sporulation efficiency of the T strain, *ert1*Δ, *pip2*Δ and *gat3*Δ significantly reduced the mean sporulation efficiency in the T strain by about 1.5-fold (*P* = 2.1 × 10^−12^, *P* = 6.1 × 10^−13^, *P* = 9.6 × 10"^10^ respectively, pair test in Methods, Figure 4C). Significant interaction terms were obtained between the genetic backgrounds (S and T) and *ert1*Δ and *pip2*Δ (*P* = 2.3 × 10^−4^, *P* = 0.04, see Methods) but not for *gat3*Δ. This showed that the effect of *ert1*Δ and *pip2*Δ on sporulation efficiency was specific to *TAO3(4477C)*, making them causal mediating genes. *GAT1* and *GAT3* were non-mediating genes, the former associated with the genotype only or a sporulation-consequential gene and the latter affected sporulation independent of the genotype. Therefore genetic and functional validation using this model identified true causal genes, namely *ERT1* and *PIP2*, mediating the effect of the allelic variant of *TAO3* on sporulation efficiency.

## DISCUSSION

Strong effects on phenotypic variation have been observed as a consequence of rare coding variants (Cohen *et al*. 2004; Cohen *et al*. 2005). Tao3 is conserved from yeast to humans (Hergovich *et al*. 2006) and had been functionally annotated solely for mitotic cell division (Du and Novick 2002; Nelson *et al*. 2003) until it was mapped for sporulation efficiency variation (Deutschbauer and Davis 2005). In this study we identify *ERT1* and *PIP2* as the *TAO3(4477C)-*-dependent mediators contributing to efficient meiosis. These genetic interactors of *TAO3(4477C)* are distinct from the mitotic interactors of *TAO3(4477G)*. In this study we identify their novel regulatory role in sporulation efficiency.

Acetate is the sole non-fermentable carbon source available to yeast during sporulation in laboratory conditions. During sporulation, this acetate gets internalized into the tricarboxylic acid (TCA) and glyoxylate cycle. Gluconeogenesis utilizes the TCA cycle intermediates and synthesizes storage carbohydrates like trehalose that is utilized during late sporulation processes (Ray and Ye 2013). Hence TCA, glyoxylate and gluconeogenic metabolic processes are crucial for sporulation to proceed since reduced flux through these pathways decreases sporulation efficiency (Aon *et al*. 1996). Moreover the genes encoding the crucial enzymes of these metabolic processes such as *PFK1, CIT1* and *CIT2* are essential for sporulation (Deutschbauer *et al*. 2002). *ERT1* and *PIP2* are known to regulate these metabolic enzymes (Baumgartner *et al*. 1999; Gasmi *et al*. 2014). Taken all together, our results suggest that the rare sporulation-associated *TAO3(4477C)* allele by interacting with regulators of TCA cycle and gluconeogenic enzymes, modulates the metabolic flux early during sporulation to result in a better sporulation efficiency.

*IME1* acts as a bottleneck for sporulation decision pathway. Lorenz and Cohen (2014) observed that many sporulation-associated natural polymorphisms have been identified in genes upstream or interacting with this input/output gene *IME1*. One of such polymorphic genes *RIM15* is a nutrient-sensing regulator of *IME2*. While *TAO3* and *MKT1* (Gupta *et al*. 2015) do not directly regulate *IME1*, we show that variants in these two genes regulate early upstream metabolic processes that impinge on *IME1*. Thus our study provides support for the hypothesis that genes surrounding the signal transduction bottlenecks are reservoirs for accumulating causal genetic variants.

During mitosis Tao3 localizes to polarized bud sites (Nelson *et al*. 2003). Further determination of co-localization of *TAO3(4477C)* with membrane-associated *ERT1* and beta-oxidation regulators *OAF1-PIP2* will give interesting clues of its function during sporulation. Similar to other scaffolding proteins like *Fry* (*Drosophila*) and *SAX-2* (*C. elegans*), Tao3 has multiple conserved Armadillo-like repeats (Hergovich *et al*. 2006) and the causal sporulation variant is present in one of them. Tao3(1493E) physically interacts with the RAM network proteins in rich growth conditions. It would be interesting to determine binding partners of Tao3(1493Q) during sporulation and if the variant affects the binding of this putative scaffolding protein. Additionally a few genes enriched for iron metabolism were differentially expressed during growth phase prior to incubation in the sporulation medium (t = 0h). It would be interesting to study whether this metabolic effect of *TAO3* also plays a role in sporulation.

Even if the basic cellular network of an organism is known, it is crucial to understand how natural genetic variation and stress conditions modulate the molecular interactions within this network resulting in differences in phenotypic outcomes (Gasch *et al*. 2016). Studies, such as this work, aimed at understanding the molecular consequences of genetic variation are especially important in the field of personalized medicine to make more reliable predictions regarding the functional consequences of an individual’s genotype on disease predisposition and treatment (Burga and Lehner 2013).

## MATERIALS AND METHODS

### Yeast strains and media

The yeast strains were grown in standard conditions at 30°C in YPD (1% Yeast extract, 2% Bacto peptone, 2% dextrose). Allele replacement strain YAD331 (Deutschbauer and Davis 2005) was a S288c-background diploid strain containing the homozygous causative sporulation polymorphism *TAO3(4477C)*. Whole-genome resequencing of YAD331 with S288c strain as the reference strain identified two additional polymorphisms (Figure S6, Table S7). Three consecutive backcrosses were performed between the haploid derivative of YAD331 and the haploid reference strain (S288c) to remove these secondary polymorphisms. After the backcrosses, the sole genetic difference between the reference S288c strain and the backcrossed allele replacement strain was at *TAO3(G4477C)* position, which was confirmed by performing PCR-based sequencing 650bp up and downstream around the two secondary polymorphisms and the *TAO3* polymorphic nucleotide. This backcrossed strain was diplodized to make it homozygous at *TAO3(4477C)* position and was termed as “T strain” in this study. The diploid parental strain S288c was termed as “S strain” in the study. All gene deletions in the study were made in the haploids of T and S strains except the ones made in SK1 strain (Table S8). Deletions were performed and verified as described previously (Goldstein and McCusker 1999; Gietz and Woods 2002). The haploid strains were diplodized using pHS2 plasmid (containing a functional *HO*) and mating type were confirmed by performing MAT PCR (Huxley *et al*. 1990). All the experiments in this study were performed using the diplodized parent strains and their diploid derivatives. For replacing the endogenous *TAO3* promoter (−150 to −1 bp upstream start site) in the T strain with a tetracycline-responsive promoter, a *tetO_7_*-based promoter substitution cassette containing *kanMX4* was amplified from the plasmid pCM225 (Bellí *et al*. 1998b). The diploid T strain with this tetO_7_-based cassette is termed P_Tet_-*TAO3(4477C)* strain. The primers for sequencing, deletions and their confirmations are listed in Table S9.

### Phenotyping

Sporulation efficiency estimation at 48h, progression through meiotic landmark events Meiosis I (MI) and Meiosis II (MII) and its quantitation was done as described previously (Gupta *et al*. 2015). For quantitation of meiotic landmarks in the T strain, parametric curves assuming delayed and 1st order kinetics were fitted to the DAPI-stained meiotic progression time course data and fitting uncertainties were estimated by bootstrapping (File S1). Cell cycle progression data for S288c and SK1 strains was taken from Gupta *et al*. (2015) (Figure 1D–E). Conditional expression of *TAO3(4477C)* was performed by constructing P_Tet_-TAO3 strain (details in File S1), which was responsive to tetracycline-analogue doxycycline (Bellí *et al*. 1998a; Bellí *et al*. 1998b). Doxycycline (2μg/ml) was added in growth and sporulation media to decrease the expression of *TAO3* gene. For each strain, a minimum of three biological replicates was used and the experiment was carried out a minimum of two times. Approximately 300 cells were counted per replicate. Fold difference was calculated as the ratio of mean sporulation efficiencies of the two strains A and B when the sporulation efficiency of A is greater than of B. Growth curve analysis was performed for individual strains grown in YPD in 96-well plates. Cells were grown overnight in YPD to saturation, reinoculated in YPD in transparent 96-well plates with a starting OD_600_ of 0.01 and grown with shaking at 30°C for 24h in Tecan Infinite M200 microplate reader. Doubling times were calculated from OD measurements of liquid cultures at a wavelength of 600 nm in the Tecan reader. For each strain, four technical replicates for each of the three biological replicates were used. Raw sporulation efficiency values are given in Table S10.

### Statistical test for calculating sporulation efficiency

For comparing sporulation efficiency, two statistical tests were used: the pair test and the interaction test. The pair test tests the null hypothesis that the two given strains (S and T) have the same sporulation efficiency.

The number *y*_*i*,*k*_ of sporulated cells (4-nuclei count) among the total number of cells *n*_*i*,*k*_ of strain *i* in replicate experiment *k* was modeled with a quasi-binomial generalized linear model using the *logit* link function and subject to a common log-odd ratio *β*_*i*_ between replicates, i.e.:

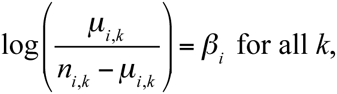

where *μ*_*i*,*k*_ = *E*(*y*_*i*,*k*_).

The pair test tests the null hypothesis of equality of log odd-ratios for two strains *i* and *j*, i.e. *H*_0_: *β*_*i*_ = *β*_*j*_

In the case of the S and T strains, the interaction test tests the null hypothesis that the effect of mutation A is independent of the effect of mutation B taking the T strain as reference background. This test thus compares four strains: mutation A only, mutation B only, both A and B and neither A nor B (T strain). Here the S strain was considered as a T strain mutated for *TAO3(4477)*. For every interaction test, we considered the dataset of the four strains of interest and fitted a quasi-binomial generalized linear model using the *logit* link function and subject to:

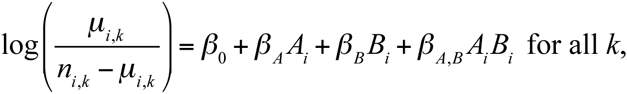

where *A*_*i*_ and *B*_*i*_ are indicator variables of the mutations A and B in strain *i* respectively. The interaction test tested the null hypothesis that the odd ratio of sporulation in the double mutant equals the product of the odd ratios of each mutation, i.e. *H*_0_: *β*_*A,B*_ = 0.

Both the pair test and the interaction test were implemented in the statistical language R with the function *glm()* assuming a constant variance function fitted by maximizing the quasilikelihood and using the t-test on the tested parameters (Gupta *et al*. 2015).

### Whole genome gene-expression profiling

Sporulating yeast cell collection at 0h, 30m, 45m, 1h10m, 1h40m, 2h30m, 3h50m, 5h40m and 8h30m (logarithmic time-series), RNA isolation and cDNA preparation were performed as previously described (Xu *et al*. 2009). Samples were hybridized to *S. cerevisiae* yeast tiling array (Affymetrix, Cat# 520055). Arrays at each time point for both the strains were normalized together using *vsn* normalization method (Huber *et al*. 2002). For qPCR, aliquots of cDNA were used in real-time PCR analyses with reagents from Kapa SYBR fast Universal qPCR master mix (Kapa Biosystems) in the Eppendorf Real-time PCR system according to manufacturer’s protocol. For each strain, four technical replicates for each of the three biological replicates were used. The primers used are given in Table S9.

### Whole genome gene-expression analysis

Within each strain, the log_2_ expression values obtained were smoothed using *locfit* at optimized bandwidth parameter *h* = 1.2 (Figure S7), base transformed for each transcript by subtracting the expression value at each time point from the baseline value at time point t = 0h (t_0_, Table S11). This log_2_ fold change value with respect to to is described as “expression” throughout the manuscript. For identifying the genes showing temporal differential expression between the T and S strains (Table S1), method implemented in EDGE software was used, which calculated statistically significant changes in expression between the T and S strains over time (Storey *et al*. 2005). The differentially expressed genes were clustered according to their temporal expression patterns using time abstraction clustering algorithm implemented in the TimeClust software (Magni *et al*. 2008, see File S1). Four major clusters were identified in each strain: Cluster I (early trend), Cluster II (increasing trend), Cluster III (late trend), Cluster IV (repressing trend) (Table S2). The transcription factors regulating a cluster of genes were extracted using the YEASTRACT database (Teixeira *et al*. 2013). Only those transcription factors were considered as candidate genes whose target genes were significantly enriched in the corresponding cluster (*P* ≤ 0.05, odds ratio ≥ 1.5). YEASTRACT database was also used to obtain the regulation matrix of yeast for identifying target genes of regulators in this study such as *UME6*. Target genes for *ACE2* were obtained from Nelson *et al*. (2003). Significantly enriched Gene Ontology terms by biological process (Bonferroni corrected *P* ≤ 0.05, Table 1) were obtained from SGD Yeastmine (Balakrishnan *et al*. 2012).

## Data availability

The array data for the T strain has been deposited in ArrayExpress (http://www.ebi.ac.uk/arrayexpress/) with accession number E-MTAB-3889. The whole genome sequence data for the T strain has been deposited in the European Nucleotide Archive (http://www.ebi.ac.uk/ena/) with the accession number PRJEB8698. The rest of the data are available as Supporting Information. The array data and the whole genome sequence data for the S strain were downloaded from Gupta *et al*. (2015). *TAO3* gene sequence data for SGRP strains (Liti *et al*. 2009) was downloaded from (http://www.moseslab.csb.utoronto.ca/sgrp/). Additional 24 *TAO3* sequences were downloaded from Saccharomyces Genome Database (SGD, http://www.yeastgenome.org/cgi-bin/FUNGI/alignment.pl?locus=YIL129C, date accessed: 01 March 2016).

## ACKNOWLEDGEMENTS

We thank Manu Tekkedil for help with whole genome sequencing sample preparation and Allan Jones and Gyan Bhanot for helpful comments. This research was supported by Tata Institute of Fundamental Research intramural funds and Department of Biotechnology grant BT/PR14842/BRB/10/881/2010 (H.S.); Bavarian Research Center for Molecular Biosystems and Bundesministerium für Bildung und Forschung through the Juniorverbund in der Systemmedizin “mitOmics” grant FKZ 01ZX1405A (J.G.); the National Institutes of Health, Deutsche Forschungsgemeinschaft and a European Research Council Advanced Investigator Grant (L.M.S.). The funders had no role in study design, data collection and analysis, decision to publish, or preparation of the manuscript.

Comparison of functional GO categories of differentially expressed genes in the T strain clusters with the S strain. See Table S2 for the full list of genes in each cluster.

Functional GO classification of the regulators of the differentially expressed genes showing early and increasing expression only in the T strain. See Tables S3 and S4 for the full list of genes.

## SUPPORTING INFORMATION

**File S1.** Detailed methods

**Table S1.** Differentially expressed genes between the T and S strains with their *P* and *Q* values calculated using EDGE

**Table S2.** Genes in each cluster using TimeClust

**Table S3.** Transcription factors regulating unique early (Cluster I) genes of the T strain

**Table S4.** Transcription factors regulating unique increasing (Cluster II) genes of the T strain

**Table S5.** Differentially expressed target genes of regulators of candidate genes mediating the affect of *TAO3*

**Table S6.** Transcription factors regulating unique repressing (Cluster IV) genes of the S strain

**Table S7.** Whole genome resequencing results for the *TAO3* allele replacement strain

**Table S8.** Strain list

**Table S9.** Primer list

**Table S10.** Raw sporulation efficiency values

**Table S11.** Smoothed expression data, base transformed with respect to t_0_ for the T and S strains

**Figure S1.** Mathematical modeling to identify stages of meiosis affected by *TAO3* causal allele

**Figure S2.** Growth phenotype and *TAO3* expression in P_Tet_-*TAO3(4477C)* strain

**Figure S3.** Comparison of global gene expression between the T and S strains at t = 0h

**Figure S4.** Comparison of genes showing early (Cluster I) and increasing trend (Cluster II) between the T and S strains

**Figure S5.** Genes having early expression in the T strain show expression at later time points or repressed in the S strain

**Figure S6.** Whole genome resequencing of *TAO3* allele replacement strain (YAD331, (Deutschbauer and Davis 2005) in comparison to the S288c reference strain

**Figure S7.** Smoothing of normalized temporal data using *locfit*

## REFERENCES

Aon, J. C., V. A. Rapisarda, and S. Cortassa, 1996 Metabolic fluxes regulate the success of sporulation in *Saccharomyces cerevisiae*. Exp Cell Res 222: 157–162.

Balakrishnan, R., J. Park, K. Karra, B. C. Hitz, G. Binkley et al., 2012 YeastMine – an integrated data warehouse for *Saccharomyces cerevisiae* data as a multipurpose tool-kit. Database (Oxford) 2012: bar062–bar062.

Baumgartner, U., B. Hamilton, M. Piskacek, H. Ruis, and H. Rottensteiner, 1999 Functional analysis of the Zn_2_Cys_6_ transcription factors Oaf1p and Pip2p - different roles in fatty acid induction of β-oxidation in *Saccharomyces cerevisiae*. J Biol Chem 274: 22208–22216.

Bellí, G., E. Garí, M. Aldea, and E. Herrero, 1998a Functional analysis of yeast essential genes using a promoter-substitution cassette and the tetracycline-regulatable dual expression system. Yeast 14: 1127–1138.

Bellí, G., E. Garí, L. Piedrafita, M. Aldea, and E. Herrero, 1998b An activator/repressor dual system allows tight tetracycline-regulated gene expression in budding yeast. Nucleic Acids Res 26: 942–947.

Bogomolnaya, L. M., R. Pathak, J. Guo, and M. Polymenis, 2006 Roles of the RAM signaling network in cell cycle progression in *Saccharomyces cerevisiae*. Curr Genet 49: 384–392.

Burga, A., and B. Lehner, 2013 Predicting phenotypic variation from genotypes, phenotypes and a combination of the two. Curr Opin Biotechnol 24: 803–809

Cirulli, E. T., and D. B. Goldstein, 2010 Uncovering the roles of rare variants in common disease through whole-genome sequencing. Nat Rev Genet 11: 415–425.

Cohen, J., A. Pertsemlidis, I. K. Kotowski, R. Graham, C.K. Garcia et al., 2005 Low LDL cholesterol in individuals of African descent resulting from frequent nonsense mutations in *PCSK9*. Nat Genet 37: 161–165.

Cohen, J. C., R. S. Kiss, A. Pertsemlidis, Y. L. Marcel, R. McPherson et al., 2004 Multiple rare alleles contribute to low plasma levels of HDL cholesterol. Science 305: 869–872.

Deutschbauer, A. M., and R. W. Davis, 2005 Quantitative trait loci mapped to singlenucleotide resolution in yeast. Nat Genet 37: 1333–1340.

Deutschbauer, A. M., R. M. Williams, A. M. Chu, and R. W. Davis, 2002 Parallel phenotypic analysis of sporulation and postgermination growth in *Saccharomyces cerevisiae*. Proc Natl Acad Sci U S A 99: 15530–15535.

Du, L. L., and P. Novick, 2002 Pag1p, a novel protein associated with protein kinase Cbk1p, is required for cell morphogenesis and proliferation in *Saccharomyces cerevisiae*. Mol Biol Cell 13: 503–514.

Gagneur, J., O. Stegle, C. Zhu, P. Jakob, M.M. Tekkedil et al., 2013 Genotype-environment interactions reveal causal pathways that mediate genetic effects on phenotype. PLoS Genet 9: e1003803.

Gasch, A. P., B. A. Payseur, and J. E. Pool, 2016 The power of natural variation for model organism biology. Trends Genet 32: 147–154.

Gasmi, N., P. E. Jacques, N. Klimova, X. Guo, A. Ricciardi et al., 2014 The switch from fermentation to respiration in *Saccharomyces cerevisiae* is regulated by the Ert1 transcriptional activator/repressor. Genetics 198: 547–560.

Gerke, J., K. Lorenz, and B. Cohen, 2009 Genetic interactions between transcription factors cause natural variation in yeast. Science 323: 498–501.

Gietz, D. R., and R. A. Woods, 2002 Transformation of yeast by lithium acetate/singlestranded carrier DNA/polyethylene glycol method. Methods Enzymol 350: 87–96.

Goldstein, A. L., and J.H. McCusker, 1999 Three new dominant drug resistance cassettes for gene disruption in *Saccharomyces cerevisiae*. Yeast 15: 1541–1553.

Granek, J. A., Ö. Kayikçi, and P. M. Magwene, 2011 Pleiotropic signaling pathways orchestrate yeast development. Curr Opin Microbiol 14: 676–681.

Gupta, S., A. Radhakrishnan, P. Raharja-Liu, G. Lin, L.M. Steinmetz et al., 2015 Temporal expression profiling identifies pathways mediating effect of causal variant on phenotype. PLoS Genetics 11: e1005195.

Gurvitz, A., F. Suomi, H. Rottensteiner, J. K. Hiltunen, and I. W. Dawes, 2009 Avoiding unscheduled transcription in shared promoters: *Saccharomyces cerevisiae* Sum1p represses the divergent gene pair *SPS18-SPS19* through a midsporulation element (MSE). FEMS Yeast Res 9: 821–831.

Hergovich, A., M. R. Stegert, D. Schmitz, and B. A. Hemmings, 2006 NDR kinases regulate essential cell processes from yeast to humans. Nat Rev Mol Cell Biol 7: 253–264.

Huber, W., A. von Heydebreck, H. Sültmann, A. Poustka, and M. Vingron, 2002 Variance stabilization applied to microarray data calibration and to the quantification of differential expression. Bioinformatics 18: S96–S104.

Huxley, C., E. D. Green, and I. Dunham, 1990 Rapid assessment of *S. cerevisiae* mating type by PCR. Trends Genet 6: 236.

Jambhekar, A., and A. Amon, 2008 Control of meiosis by respiration. Current Biol 18: 969–975.

Karpichev, I. V., and G. M. Small, 1998 Global regulatory functions of Oaf1p and Pip2p (Oaf2p), transcription factors that regulate genes encoding peroxisomal proteins in *Saccharomyces cerevisiae*. Mol Cell Biol 18: 6560–6570.

Kassir, Y., N. Adir, E. Boger-Nadjar, N. G. Raviv, I. Rubin-Bejerano et al., 2003 Transcriptional regulation of meiosis in budding yeast. Int Rev Cytol 224: 111–171.

Lardenois, A., E. Becker, T. Walther, M. J. Law, B. Xie et al., 2015 Global alterations of the transcriptional landscape during yeast growth and development in the absence of Ume6-dependent chromatin modification. Mol Genet Genomics 290: 2031–2046.

Lee, J. H., M. Huynh, J. L. Silhavy, S. Kim, T. Dixon-Salazar et al., 2012 *De novo* somatic mutations in components of the PI3K-AKT3-mTOR pathway cause hemimegalencephaly. Nat Genet 44: 941–945.

Lin, C. P., C. Kim, S. O. Smith, and A. M. Neiman, 2013 A highly redundant gene network controls assembly of the outer spore wall in *S. cerevisiae*. PLoS Genetics 9: e1003700.

Liti, G., D. M. Carter, A. M. Moses, J. Warringer, L. Parts et al., 2009 Population genomics of domestic and wild yeasts. Nature 458: 337–341.

Ljungdahl, P. O., and B. Daignan-Fornier, 2012 Regulation of amino acid, nucleotide, and phosphate metabolism in *Saccharomyces cerevisiae*. Genetics 190: 885–929.

Lorenz, K., and B. A. Cohen, 2014 Causal variation in yeast sporulation tends to reside in a pathway bottleneck. PLoS Genet 10: e1004634.

Magni, P., F. Ferrazzi, L. Sacchi, and R. Bellazzi, 2008 TimeClust: a clustering tool for gene expression time series. Bioinformatics 24: 430–432.

Manolio, T. A., F. S. Collins, N. J. Cox, D. B. Goldstein, L. A. Hindorff et al., 2009 Finding the missing heritability of complex diseases. Nature 461: 747–753.

Marzluf, G. A., 1997 Genetic regulation of nitrogen metabolism in the fungi. Microbiol Mol Biol Rev 61: 17–32.

Neiman, A. M., 2005 Ascospore Formation in the yeast *Saccharomyces cerevisiae*. Microbiol Mol Biol Rev 69: 565–584.

Neiman, A. M., 2011 Sporulation in the budding yeast *Saccharomyces cerevisiae*. Genetics 189: 737–765.

Nelson, B., C. Kurischko, J. Horecka, M. Mody, P. Nair, et al., 2003 RAM: A conserved signaling network that regulates Ace2p transcriptional activity and polarized morphogenesis. Mol Biol Cell 14: 3782–3803.

Primig, M., R. M. Williams, E. A. Winzeler, G. G. Tevzadze, A. R. Conway, et al., 2000 The core meiotic transcriptome in budding yeasts. Nat Genet 26: 415–423.

Ray, D., and P. Ye, 2013 Characterization of the metabolic requirements in yeast meiosis. PLoS ONE 8: e63707.

Saint Pierre, A., and E. Génin, 2014 How important are rare variants in common disease? Brief Funct Genomics 13: 353–361.

Spellman, P. T., G. Sherlock, M. Q. Zhang, V. R. Iyer, K. Anders et al., 1998 Comprehensive identification of cell cycle-regulated genes of the yeast *Saccharomyces cerevisiae* by microarray hybridization. Mol Biol Cell 9: 3273–3297.

Stankiewicz, P., and J. R. Lupski, 2010 Structural variation in the human genome and its role in disease. Annu Rev Med 61: 437–455.

Storey, J. D., W. Xiao, J. T. Leek, R. G. Tompkins, and R. W. Davis, 2005 Significance analysis of time course microarray experiments. Proc Natl Acad Sci U S A 102: 12837–12842.

Teixeira, M. C., P. T. Monteiro, J. F. Guerreiro, J.P. Gonçalves, N.P. Mira et al., 2013 The YEASTRACT database: an upgraded information system for the analysis of gene and genomic transcription regulation in *Saccharomyces cerevisiae*. Nucleic Acids Res 42: D161–D166.

Tomar, P., A. Bhatia, S. Ramdas, L. Diao, G. Bhanot et al., 2013 Sporulation genes associated with sporulation efficiency in natural isolates of yeast. PLoS ONE 8: e69765.

Turcotte, B., X. B. Liang, F. Robert, and N. Soontorngun, 2010 Transcriptional regulation of nonfermentable carbon utilization in budding yeast. FEMS Yeast Res 10: 2–13.

Voth, W. P., A. E. Olsen, M. Sbia, K. H. Freedman, and D. J. Stillman, 2005 *ACE2, CBK1*, and *BUD4* in budding and cell separation. Eukaryotic Cell 4: 1018–1028.

Xu, Z., W. Wei, J. Gagneur, F. Perocchi, S. Clauder-Münster et al., 2009 Bidirectional promoters generate pervasive transcription in yeast. Nature 457: 1033–1037.

Zuk, O., S. F. Schaffner, K. Samocha, R. Do, E. Hechter et al., 2014 Searching for missing heritability: designing rare variant association studies. Proc Natl Acad Sci U S A 111: E455–E464.

